# A 96-WELL VALVED MICROFLUIDIC DEVICE FOR TESTING OF LIVE INTACT TUMOR CUBOIDS

**DOI:** 10.1101/2022.07.07.499178

**Authors:** Ethan J. Lockhart, Lisa F. Horowitz, Cb Lim, Tran Nguyen, Mehdi Mehrabi, Taranjit S. Gujral, Albert Folch

## Abstract

There is a pressing need for functional testing platforms that use human, live tumor tissue to better predict traditional and immunotherapy responses. Such platforms should also retain as much of the native tumor microenvironment (TME) as possible, as many cancer drug actions rely on TME-dependent mechanisms. Present high-throughput testing platforms that have some of these features, e.g. based on patient-derived tumor organoids, require a growth step that alters the TME. On the other hand, micro-dissected tumor tissue “spheroids” that retain an intact TME have shown promising responses to immunomodulators acting on native immune cells. Here we demonstrate a microfluidic 96-well platform designed for drug treatment of hundreds of similarly-sized, cuboidal micro-tissues (“cuboids”) produced from a single tumor sample. Four cuboids per well are automatically arrayed into the platform using hydrodynamic trapping. The microfluidic device, entirely fabricated in thermoplastics, features microvalves that fluidically isolate each well after the cuboid loading step. Since the platform effectively makes the most of scarce tumor tissue, we believe it could ultimately be applied to human biopsies for drug discovery and personalized oncology, altogether bypassing animal testing.

## INTRODUCTION

Drug discovery and development are vastly inefficient. For more than a century, biologists have performed pre-clinical drug tests in Petri dishes and animal models. One of the limitations of such models is that they lack the original human tumor microenvironment (TME).^1^ As a result, drugs that have passed pre-clinical tests are not necessarily effective in the context of the human TME. Only ~14% of the ~1,000 drugs in clinical trials each year pass the safety and efficacy tests, a number that reaches a dismal <4% for cancer drugs;^2^ more than half of the failures are due to lack of efficacy.^3^ Due to this inefficiency, the average drug takes >10 years and >$1 billion to come to market.^4^ Most critically, the inefficiency of drug development has an impact far beyond the pharmaceutical industry. On average, pharma needs to recover the money invested in the failed drugs by charging the losses to the approved drugs, and these are the drugs ultimately billed to hospitals and patients. Also, the long periods spent in drug development means more time the disease put avoidable strain on healthcare and society.

In the last decade, functional drug testing technologies such as patient-derived organoids^5^ and organs-on-chips^6^ have emerged as promising new approaches. However, in these platforms, which often take weeks to months to establish, the tumor tissue is generated *de novo* from the patient’s cancer cells^7^ and often utilize non-native extracellular matrix. Therefore, these *ex vivo* models do not preserve the patient’s original, unique TME and suffer from *in vitro* cultivation bias. A different paradigm for cancer drug development that preserves the human TME is critically needed to help meet the pace of emerging new therapies while keeping treatments more affordable.

Functional assays on intact tissues can potentially complement and extend genomics-based and/or organoid-based approaches for drug testing by capturing key determinants of therapeutic response such as tissue architecture, tumor heterogeneity, and the TME.^8^ Researchers have developed diverse functional assay platforms that assess drug responses in tumor samples (slices, spheroids, etc.).^9,10^ In recent years, patient-derived tumor “organoids” have shown great promise to predict drug responses for personalized cancer treatment^11–14^. Also termed “tumor spheroids”, these are produced by dissociation of a cancer patient’s live biopsy and subsequent growth of the cells in a 3D architecture. Tumor spheroids are thought to more faithfully replicate the drug responses seen *in vivo* than simple 2D culture because they recapitulate some of the cellular and molecular relationships present in the TME.^15^ Critically, however, the growth steps to create and expand tumor spheroids come at the expense of loss of much of the native immune cells and their original 3D relationships within the TME – limiting the relevance of tumor organoids, and especially as models for immunotherapy. Many immunotherapy drugs act on the local TME.^16–18^ Micro-dissected tumors (“μDTs”), also termed “*ex vivo* tumor fragments”, are a spheroid approach that has been used for decades to preserve the tumor and its TME (with its native immune cells) within manually-cut small (~1-3 mm-wide) pieces.^19–23^ μDTs have recently shown promise for study of immunotherapy with checkpoint inhibitors, but at very low throughput.^24–29^ The μDTs were minced manually with a scalpel and grown in collagen gel either on Transwells^27–29^ or in microchannels.^25,26^ Astolfi et al. prepared ~420 μm-diam. cylindrical μDTs by punching cores from PDX tumor slices and manually loaded them in microchannels with 5 traps each; the reproducible size of the μDTs allowed for a metabolite transport model.^30^ Implantable or needle microdelivery devices^31,32^ locally deliver small doses of (up to 16) drugs to the tumor *in vivo,* with maximal preservation of the TME, but issues of tumor accessibility and patient safety limit their applicability. PDX mouse models permit study of drug responses in an intact organism (including immune checkpoint blockade in humanized PDX).^33^ Champions Oncology now offers a screen using *ex vivo* tumor fragments that contains 32 EGFR+ HER2+ PDX models across a wide range of cancers (breast, pancreatic, colorectal, etc.). However, PDXs have the critical disadvantage that all or most of the TME is from the host mouse, and PDX from individual patients grow too slowly to inform initial post-operative therapeutic decisions.

To address the above limitations, we have recently developed a microfluidic platform for drug development and precision medicine based on regularly-sized, cuboidal-shaped microdissected tissues (referred to as “cuboids”) that are mechanically cut.^34^ In min, more than 10,000 cuboids (~400 μm-wide) can be produced from ~1 cm^3^ of solid tumor. The cuboids are never dissociated and retain much of the native TME (e.g. immune cells and microvasculature). Our initial platform used differential hydrodynamic flows to trap and array 24 cuboids with 3 cuboids per each of 8 wells.^34^ Here, we present a different design in which our user-friendly multi-well platform can hydrodynamically trap and selectively treat arrays of cuboids (96 wells, 384 cuboid traps, with 4 traps per well), with pressure-controlled TPU valves to prevent cross-contamination between wells.

## METHODS

### Computer-aided design of the microfluidic device

To facilitate design iteration, we used the 3D CAD software, Autodesk Inventor, to model the layers of the device. Our 96-well microfluidic platform consisted of 5 operational layers, with an additional bottomless well plate intended for attachment at the end, during the experiment after cuboid loading and before culture. For the microchannel layer (MCL) the main fluidic channels were 700 μm in depth and 400 μm in width to minimize fluidic resistance. We made the trap dimensions 500 μm in depth and 800 μm in diameter, just large enough to hold 1 cuboid. To prevent cuboids from entering the microchannel network through the traps, we made the channels that connect the traps to the main fluidic channels shallower (100 μm). For the valve seat layer (VSL), we made the depth of the valve seat 50 μm to minimize the travel distance required for the valving component to achieve closure via pneumatic control. The width of the valve seat was 390 μm. We included square through-holes on each side of the valve seat to enable fluidic communication with the MCL once the two layers were bonded. For the pneumatic control layer (PCL), the depth of the channels was 300 μm and the width was 200 μm. These dimensions minimized the potential for the valve membrane material, thermoplastic polyurethane (TPU), to bond to the inside walls of the channels during the manufacturing process. The circular chambers were 2 mm in diameter, and we positioned these to align with the rest of the valve architecture located in the other layers. Additionally, we modeled both the bottomless well plate and loading frame to match the overall size of a 96-well plate. Finally, we imported all 3D models to Fusion 360 (Autodesk).

We used the built-in computer-aided manufacturing (CAM) tools within Fusion 360 to generate toolpaths for our computer numerical control (CNC) milling machine. The 2D-pocket CAM function, which generates toolpaths based on selected areas of a 3D model, created toolpaths for microchannel and through-hole features. However, due to the size and complexity of our device, we avoided selecting entire networks of channels within single 2D-pocket functions which would otherwise crash the program. Instead, we segmented the large channel networks into loadable portions using sketch tools. Closed 2D sketch contours also worked to designate milling areas. After dividing the channel network into smaller portions using sketch outlines, we could repeat the 2D-pocket operation and select manageable portions of the network within each instance. To generate toolpaths for all other geometries, we utilized the 2D-contour and 3D-parallel functions. In particular, the 3D-parallel operation allowed us to create the toolpath that cut the smooth curvature of the valve seats in the VSL.

### CNC milling device components in PMMA

To fabricate the device, we used a 3-axis CNC milling machine (DATRON neo) to cut features on polymethyl methacrylate sheets (PMMA, Astra Products Inc.). Our 96-well microfluidic platform consisted of 5 layers. The primary 0.5 mm-thick MCL was cut using a 381 μm diameter square endmill with a spindle speed (SS) of 40,000 rpm and a cutting feed rate (FR) of 800 mm/min. These milling parameters struck a balance between minimizing wear on the microscale endmill and completing the cut within a reasonable amount of time (≈1 hour). For the larger features of the MCL such as the 800 μm-diameter trap through-holes, we used a 1/32”diameter square endmill (SS = 40,000 rpm, FR = 850 mm/min). Note that the thickness of individual PMMA sheets varied in random areas by approximately 20% of the stock thickness due to the cell casting process of the manufacturer. Thus, calibrating the mill at one height was not reflective of the whole material surface and caused variations in the depth of cut. Therefore, prior to initiating each milling operation, we used the pressure sensing calibration probe on the DATRON to measure the z-heights of 500 evenly spaced points within the cutting area of the device. Then, the CNC machine software used these points to approximate a surface profile of the plastic sheet and automatically compensate for unevenness. Therefore, the calibration profile prevented these inconsistencies from leading to manufacturing errors and hindering device performance.

We used similar methods when cutting the other layers. For the 0.5 mm-thick VSL, we cut the curved valve seats using a 381 μm diameter ball endmill (SS= 36,000 rpm, FR = 650 mm/min). Here, a ball endmill combined with the 3D-parallel CAM function achieved a valve seat concavity greater than the radius of the ball endmill (405 μm versus 381 μm). This step was necessary to achieve the desired valve seat depth (50 μm) and width (390 μm). Then, we used a 175 μm diameter square endmill (SS = 36,000 rpm, FR = 250 mm/min) to create the through-holes bookending each valve seat. Since the height of the valve seats greatly influenced valving performance, we built the surface profile using 800 points to mitigate variation. For the 0.5 mm-thick PCL, we used a 200 μm diameter square endmill (SS = 36,000 rpm, FR = 300 mm/min) to cut the channel network in quadrants according to the method described earlier. Note, the depth of these channels was greater than the width, so we used a slower FR to reduce load on the mill. We also alternated the milling coolant to switch between ethanol and air between each quadrant operation to avoid PMMA overheating and sticking to the tip, especially during deeper passes. We measured the surface profile using 450 points since variations in channel depth were less impactful, and fewer points reduced measuring times for each layer.

For both the 6.35 mm-thick PMMA (1227T569, McMaster-Carr, Elmhurst, IL) loading frame and bottomless well plate, we lined one face of the sheet with 50 μm-thick 3M™ High-Strength Acrylic Adhesive 300LSE then used a CO2 laser system (VLS3.60, Scottsdale, USA) for fabrication. We preferred to laser cut these components because the feature dimensions exceeded 500 μm, and the acrylic adhesive layer was not amenable to CNC milling though it was necessary for device assembly.

### Device assembly

#### Cleaning CNC-milled components

Immediately following CNC milling, we removed burrs and plastic debris from the substrates through a series of washing steps. For each layer, we first rubbed liquid dish soap on the surface with a gloved hand while applying pressure. Then we rinsed the layers under water, dried them using an air gun, and inspected them under a microscope. We removed remaining burrs by reapplying soap and scrubbing with a toothbrush, angling the head to ensure the bristles reached into channels and through-hole features. After drying with air, we stored layers in a covered container.

#### Bonding PMMA layers

After cleaning, we bonded the MCL and VSL using our established thermal solvent bonding process.^34^ Briefly, PMMA becomes slightly adhesive when exposed to chloroform vapor, causing polymer reflow.^35,36^ An irreversible cohesive bond forms when two exposed surfaces are pressed together allowing us to assemble PMMA layers. As an additional benefit, treatment with chloroform vapors also reduces surface roughness caused by mill marks, improving optical quality. We filled a glass container with 50 mL of chloroform and placed four steel standoffs (6 mm) in the container that provided a 3 mm elevation from the surface of the liquid. We first exposed the bottom (open channel side) of the MCL for 2 min by setting the corners of the layer down on the standoffs. Then, we transferred this layer from the container onto an alignment tool with the exposed side facing up. The alignment tool consisted of metal pins (5 mm diameter x 15 mm length) inserted into four holes milled into the corners of a block of PMMA. These pins corresponded to the four corner holes included on each layer of the device that allowed us to consistently position layers on top of each other during bonding. After replenishing the chloroform in the container to 50 mL, we exposed the top (featureless side) of the VSL for 4 min. Then, we immediately (within 15 seconds) transferred the VSL exposed-face-down onto the MCL on the alignment tool, and hand-pressed the layers together to form a weak bond. To increase bonding strength, we sandwiched the layers between two 3 mm-thick PDMS slabs sharing the same outer dimensions as the device and compressed the stack of layers in a heat press for 5 min at 140 °F and 420 psi.

#### Bonding TPU

We incorporated valve architecture using a bonding process^53^ that utilizes temperature, pressure, and time to join a thermoplastic polyurethane film (TPU, PT9200US NAT, Covestro LLC, USA) and the pneumatic control layer (PCL) to the bottom (featured side) of the VSL simultaneously. Most of the steps of this process (which we refer to as “thermal vacuum bonding”) were necessary to remove entrapped chemicals, solvents, and gas bubbles within the bulk of the thermoplastic materials. Otherwise, gas bubbles released during the heating process likely prevented bonding in some areas or caused deflection that resulted in the TPU film sticking to the inner walls of PMMA channels rather than lay flat across them as intended (a problem we referred to as “inner-channel bonding”).

To circumvent these problems, we first performed a treatment step to prepare the PMMA materials. First, we cleaned the PMMA layers with soap as above, taking special care to remove water trapped inside the mostly-closed channels of the bonded MCL and VSL layers by aiming the air gun directly into inlets, traps, and open through-holes near the valve seats. Then we used a lint-free wipe soaked in ethanol to thoroughly scrub the surface of each layer. Following this cleaning process, we thermally treated the layers under vacuum by placing them in a vacuum oven for at least 8 hrs at 80 °C and −12psi.

Afterwards, we repeated the above cleaning process for the PMMA layers to further remove any released solvents or chemicals. We cut the TPU film with slightly larger outer dimensions than that of the device layers and wiped both sides using the ethanol-soaked lint-free wipe. As, TPU film curls into itself upon ethanol exposure, we used pressurized air to dry and flatten the TPU. For thermal treatment, we placed all layers including the TPU film inside the vacuum oven for 2 hrs at 50 °C and −12 psi. This second treatment step further removed gases and solvents that may have dissolved into the thermoplastic materials during cleaning^37^. When finished, we painted the valve seats on the VSL with a 1:1 v/v dilution of liquid dish soap and water using a ball of cotton. To produce functional valves, TPU must be prevented from bonding to the valve seats. Normally, the TPU film laid flat over sufficiently deep channels, but the likelihood of “inner-channel bonding” increased when the valve seat was shallow (depth of 50 μm). Liquid dish soap provided a temporary coating, which prevented irreversible adhesion of the TPU to the PMMA valve seat during the final thermal vacuum bonding step.

To create a temporary bond and eliminate wrinkles, we laminated the TPU film onto the surface of the VSL using a laminator (Sky, 325R6) set to 110 °C. Here we used a scalpel to cut TPU away from blocking the control inlet on the VSL. Then, we aligned these layers to the PCL. First, we placed the PCL with the channel side facing up onto the alignment frame as previously described, then we set the laminated VSL with the TPU facing down onto the pins of the frame. The laminated TPU frame covered the alignment holes, which prevented the layer from sliding down along the length of the pins, so we flipped the apparatus to bring the PCL down to contact the TPU surface. Once aligned, we sandwiched the ensemble between two 25 cm x 12.5 cm large glass slides, then used 6 paper binder clamps (3 on each side) to apply pressure. For the final bonding step, we placed the sandwich in the vacuum oven for 1 hour at 130 °C and −12 psi, then allowed the device to cool by sitting at room temperature for at least 40 min. After bonding, we ran water through the microchannels, which washed away the soap and unstuck the valves.

#### Attaching the loading frame

To create a fluid reservoir on the top of the device, we attached the PMMA loading frame to the top of the MCL via pressure sensitive adhesive (3M™ 300LSE) and pressed for 5 min at 160 psi. Finally, we inserted connection tubing into the 5 inlet holes of the loading frame and secured them by applying ethyl cyanoacrylate glue (Krazy^®^ Glue, North Carolina, USA) around the top perimeter of the holes and waiting overnight.

### Cuboid preparation

We created thousands of cuboids following our microdissection procedure as previously described.^34^ Briefly, we affixed fresh tumor tissue onto a PMMA disk using cyanoacrylate glue, and used a mechanical tissue chopper (McIlwain-type) to create 400 μm-thick slices. We then laid slices on their sides to be cut by the chopper into thin strips of tissue. Finally, we rotated the glass slide 90 degrees to be cut by the chopper into cuboids. Cuboids were then size-selected for passage through a 750 μm filter and not through a 300 μm filter (Pluriselect, USA).

### Cuboid loading

To prepare the device for loading with live tumor tissue, we first filled the microchannel network with phosphate buffered saline (PBS) using a handheld syringe and positive pressure. We used a 200 μL pipette to help remove any air bubbles created in the traps. We aspirated PBS from the reservoir, leaving enough volume to still cover the traps. We then filled the reservoir with 20% w/w polyethylene glycol (PEG, 8k M.W.) in PBS. To control fluidic suction, we connected one of the four fluidic quadrant outlets to a 60 mL syringe (BD Bioscience, San Jose, CA) and syringe pump (Fusion 200, Chemyx Inc., Stafford, TX). To control the valves, we connected the pneumatic control outlet to a microfluidic flow controller (OB1 MK3+, Elveflow, Paris, FR). We used a smaller PMMA frame (2 cm x 2 cm x 6.35 mm, fabricated via laser cutting) as the “loading window”. The “loading window” confined cuboids within a smaller area of the total reservoir, allowing us to promote hydrodynamic capture by sliding multiple cuboids over each trap. We placed the window inside the reservoir and deposited cuboids in its center.

To create flow in the microchannel network for the capturing process, we engaged suction by pulling with the syringe pump at a flow rate of 200 mL/hr from one of the four quadrants of the device. To ensure wide open valves, we applied negative pressure (−4 psi) to the single pressure control inlet during the loading process. We slid the loading window along the traps contained within the activated quadrant network until the traps were filled, then deactivated suction. We used tweezers to help redistribute the cuboids around. We continued filling the rest of the device by reconnecting the syringe pump to the control inlet of the next microchannel quadrant and repeating the process. We replenished the supply of PEG/PBS by directly pouring fresh solution into the reservoir as needed (approximately 80 mL total). Following loading, we aspirated fluid from the surface of the device, with one rinse with PBS, then attached the bottomless 96-well plate on top of the MCL using pressure-sensitive adhesive. To culture the cuboids for subsequent assays, we filled the wells with DMEM/F12 (Invitrogen) cell culture media supplemented with 5% heat-inactivated fetal bovine serum (VWR) and penicillin/streptomycin (Invitrogen).

### Fluorescent staining and imaging cuboids

Prior to staining, we closed the valves of the device by applying +4 psi to the pneumatic control inlet. Then, we stained the live cuboids for 1 hour at 37 °C with the following dyes diluted in culture medium: Cell Tracker Green CMFDA (CTG; Invitrogen, 5 μM), Cell Tracker Orange CMRA (CTO; Invitrogen, 5 μM), and Hoechst (H; Invitrogen, 16 μM). We then replaced the staining solution with live cell culture media for imaging. We imaged the device with an epifluorescence microscope (BZ-X800, Keyence, JP) using a 2x objective lens. Last, we used FIJI to adjust brightness and contrast to increase cuboid visibility.

## RESULTS/DISCUSSION

### Cuboid microdissection

We created thousands of uniformly sized, cuboidal-shaped micro-dissected tissues (“cuboids”) of (400 μm)^3^ size from Py8118 syngeneic mouse breast cancer as reported previously.^34^ We performed three series of orthogonal cuts using a mechanical tissue chopper followed by size-exclusion filtering to maximize uniformity. Aside from being amenable to hydrodynamic trapping, this size of tissue is biologically advantageous for several reasons. For one, this size is balanced since too large of tissue loses nutrient delivery and viability leading to necrotic core development, while too small of tissue fails to preserve the tumor microenvironment (TME).^34,38^ Additionally, previous organoid^34,39^ and μDT^34,40^ models from various types of solid tumors have been of similar size (50-800 μm).

### Microfluidic device concept

The size and shape of cuboids make them amenable to hydrodynamic trapping. To demonstrate this concept, we devised a microfluidic network in which four replicates of bifurcating channels propagate suction equally to a total of 384 cuboid traps (Figure 1a). Within each quadrant, the channel network transmits suction from a single inlet to 24 units of four-trap clusters that correspond to wells of a 96-well plate, resulting in four tissue replicates per well. Note that, in this design, the wells are fluidically connected. When the wells (placed on top afterwards) are filled, changes in hydrostatic pressure brought on by moving or tilting the device could create flow from well to well, which would defeat the purpose of separating solutions in different wells. To isolate the wells, we designed valves before each well to prevent hydrostatic pressure-induced flow. The design of pneumatic valves and their control network allows us to control actuation from a single source, transmitting air pressure through a secondary network of bifurcating channels to all 96 valves simultaneously (Figure 1b&c). With this device, cuboids may be arrayed hydrodynamically by using suction to pull them into traps (Figure 1d). Each trap is sized to fit one cuboid, and channel entrances within each trap have 100 μm openings to prevent tissue from entering the fluidic network. To initiate this process, the traps are submerged in sufficiently viscous fluid, then suction from the traps generates flow which draws in nearby suspended cuboids for capture. The cuboids are suspended in a viscous 20% PEG solution which aids in cuboid movement and trapping. The hydrodynamic trapping of cuboids works best when the solution has more viscosity to transmit its movement to the cuboids. We have observed that plain cell culture medium does not offer enough viscous drag, allowing the cuboids to settle too quickly. Therefore, we use 20% w/w PEG (8k M.W.) in PBS to achieve high-density, high-viscosity solutions for seeding cuboids^41^. Note that with one valve per well, any two traps are separated by two valves, which we may close to avoid cross-talk between solutions in the wells.

**Figure 1:**
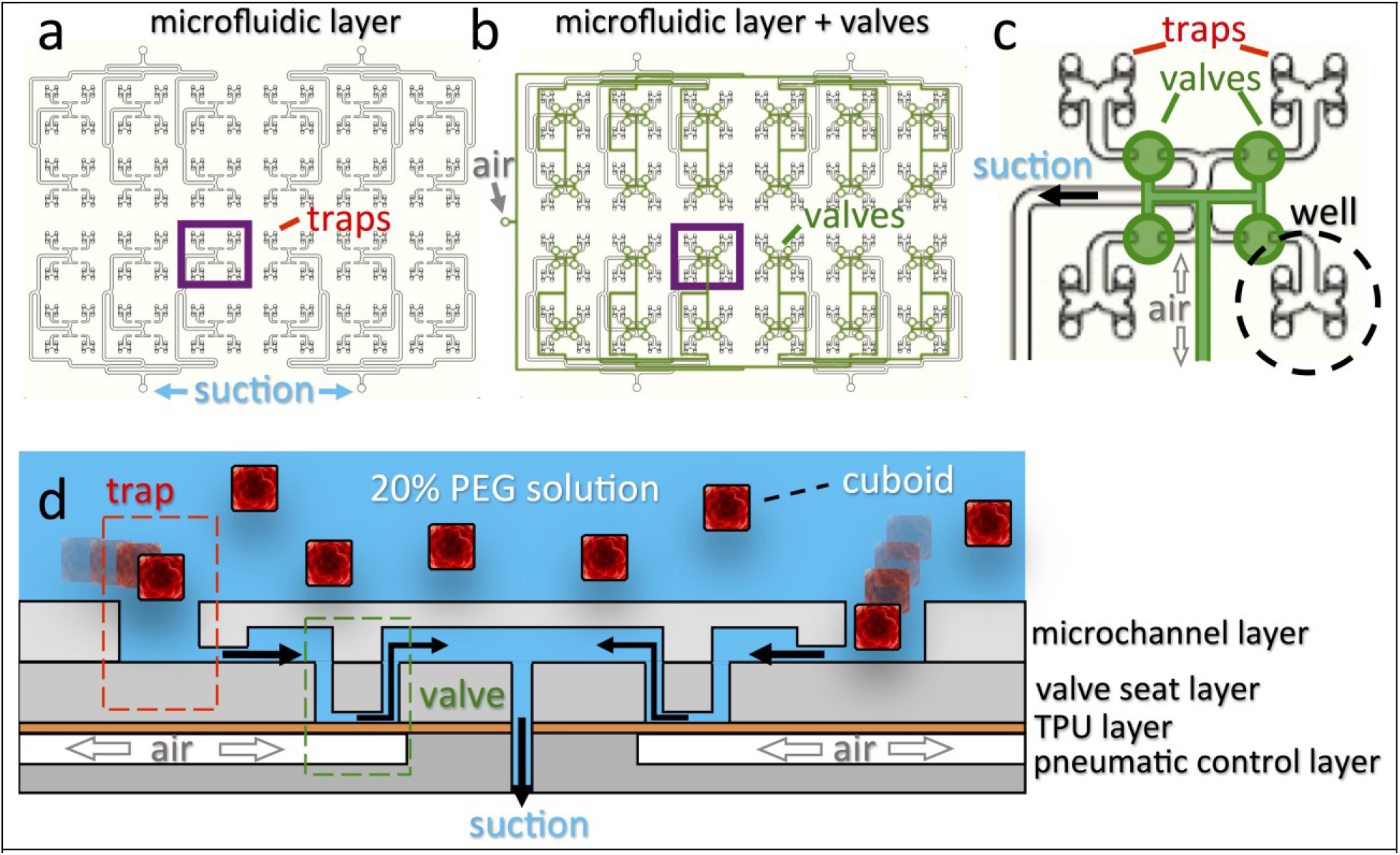
Microfluidic device concept design for hydrodynamic capture of cuboids. (a) CAD drawing of the microchannel network consisting of four fluidic quadrants and 384 traps. In each quadrant, suction propagates from a single inlet to 24 units of four-trap clusters, which are eventually separated into wells. (b) Overlay of the pneumatic control network (green) on top of the microfluidic network showing the propagation of pressure from a single outlet (gray, far left) to 96 individual valves. Two thermoplastic layers (the PMMA valve seat layer and TPU film) separate the overlaid networks as part of the valve architecture. (c) Close-up of the networks underlying 4 wells, showing separation by one valve each. Each well contains 4 tissue replicates. (d) Cross-section schematic of the device showing the cuboid-capturing process. Cuboids are suspended in a reservoir of viscous 20% PEG solution above the traps (red) and are pulled into traps via hydrodynamic suction. Following capture, positive air pressure (gray) actuates the valves to prevent well crosstalk through the fluidic channels by deflecting TPU upward into the valve seat layer.

### Device fabrication and assembly

To implement this design, we selected PMMA as the primary material for our device because it has beneficial properties for both research and clinical settings. These traits include low-cost, optical clarity, and biocompatibility, which are paramount for biological studies^42^. Additionally, PMMA avoids the drawbacks of polydimethylsiloxane (PDMS), one of the most common materials used for microfluidic device manufacturing. PDMS suffers from both molecular compound absorption^43–50^ into and adsorption^41^ onto the material. Such problems can affect the outcomes of drug-based studies by changing target concentrations or sequestering molecules in undesired regions of the device. Thus, PMMA is a more appropriate material for fabricating microfluidic drug testing platforms. In terms of manufacturability, PMMA is a rigid thermoplastic and thus cannot be molded without access to expensive equipment. However, the rigidity of the material is suitable for rapid prototyping via CNC milling – a method that provides precise control over fluidic geometries and produces a quality surface finish. Furthermore, thermoplastics can translate to high-volume manufacturing methods such as hot-embossing and injection molding which can facilitate commercial translation. However, our device also requires microvalve integration for fluidic isolation (see below). Discarding PDMS poses an obvious challenge for the integration of microvalves in a PMMA device, as a wide range of PDMS actuators can easily be molded by soft lithography^51–56^ but these cannot be straightforwardly fabricated in PMMA. Our design incorporates a microvalve where the flexible layer is made of thermoplastic urethane (TPU), adapted from a design by Shrike Zhang’s lab.^37^ Contrary to the Zhang design, our microvalve is open at rest, requiring pressure to close it.

We fabricated the 5-layered device (Figure 2a) in thermoplastic sheets via CNC milling and 3 assembly steps, utilizing thermal solvent bonding, thermal vacuum bonding, and pressure sensitive adhesive. First, we designed the loading frame, microchannel layer (MCL), valve seat layer (VSL), and pneumatic control layer (PCL) in Autodesk Inventor CAD software. The MCL contains 4 identical quadrants of a bifurcating channel network that propagates suction equally to an array of traps in a 96-well plate arrangement. The remaining 3 layers below the MCL form the architecture of the microvalves that prevent well-to-well crosstalk through the underlying channels. The loading frame atop the MCL creates a fluid reservoir. The resulting device requires two types of control lines for operation: one for each quadrant that engages fluidic suction through the microfluidic network in order to attract cuboids towards traps, and one that actuates all 96 microvalves in unison through the pneumatic control network (Figure 2b). We made the channels of the MCL 700 μm wide and 400 μm deep to minimize fluidic resistance and increase suction from the traps, while the channels of the PCL were 200 μm wide and 300 μm deep to prevent TPU from collapsing and bonding with the inside walls of the channels during the thermal vacuum bonding process.

**Figure 2:**
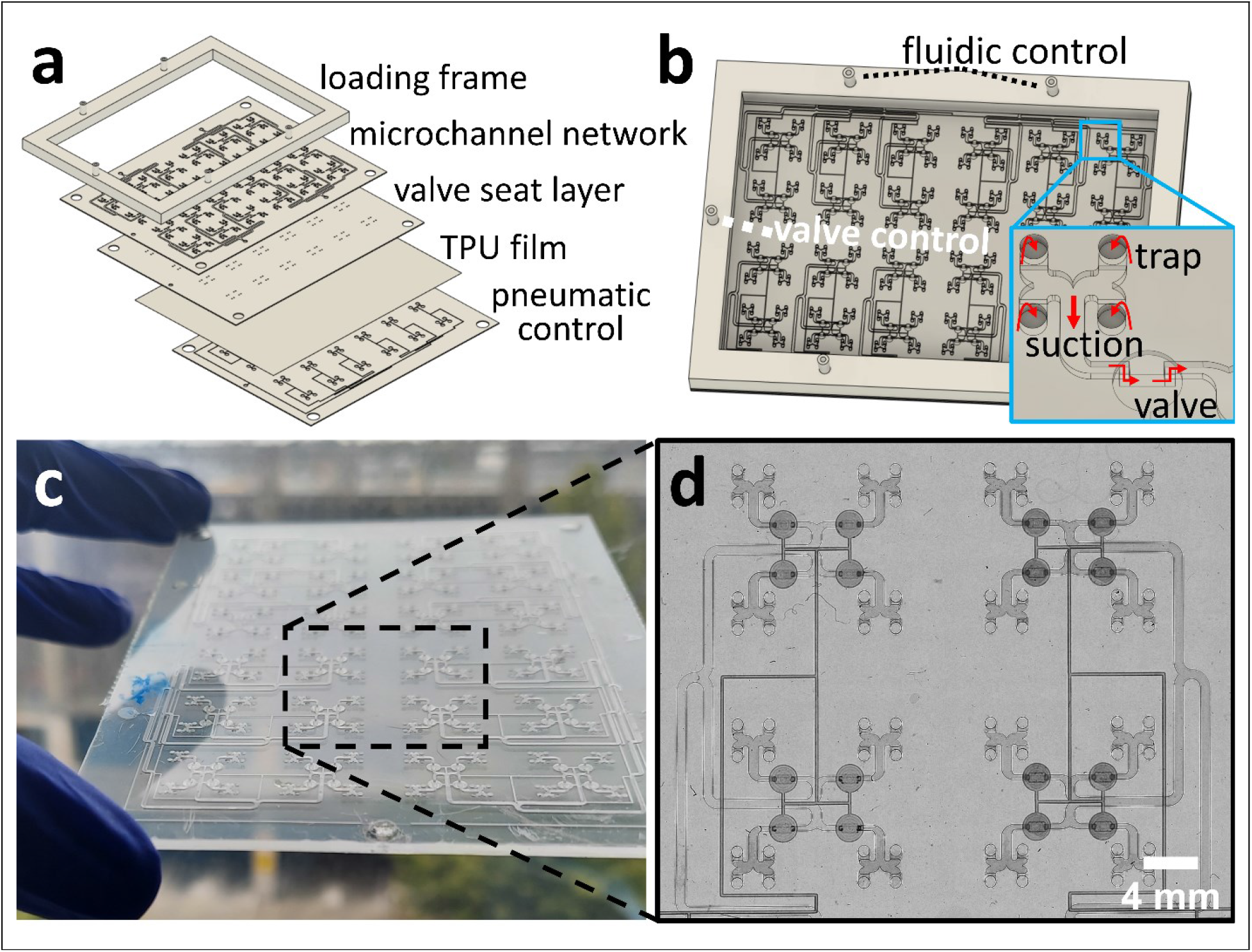
Assembly of the microfluidic device. (a) 3D CAD exploded view showing the 5 functional device layers. (b) Perspective view of the assembled device with inset showing the direction of flow. (c) The assembled layers prior to addition of the loading frame. (d) Top view close-up of 16 wells showing traps and valves of the fully-bonded chip.

Next, we manufactured each layer using CNC milling. The MCL and PCL were cut using square endmills, while the VSL, which consisted of an array of concave channels and connecting through-holes, was cut using a ball endmill for the concave portions and a square end mill for the through-holes. For all layers, we selected larger endmill sizes as needed to minimize cutting time and reduce wear on smaller mills, which were more prone to breakage over long cutting times (for further details see the Methods section).

Critically for work with microfluidics, stock sheets of PMMA had significant variations in thickness (approximately 20% of the stock thickness) that could potentially impact device performance. These random, unintended features can cause errors in mill calibration and introduce unintended variability (e.g. 70 μm differences in height for microchannels cut into 300 μm-thick PMMA). The plastic manufacturer’s cell casting process introduces these variations, making them unavoidable. To overcome this challenge, we utilized a function of the DATRON neo CNC mill that measures the surface profile of the material before each cut with its built-in pressure-sensing probe, also used to calibrate the z-height of the spindle. The probe can acquire up to 2,000 measurements of height evenly distributed across the surface of the workpiece that the machine then uses to extrapolate a surface profile. The CNC mill then automatically adjusts the spindle height to compensate for areas of measured unevenness. The accuracy of the extrapolated surface profile increases with the number of points measured; however, this process also prolongs manufacturing.

After milling, we assembled the device. We first removed burrs and debris, then used our established thermal solvent bonding technique ^34^ to bond the bottom (open channel side) of the MCL to the top (featureless side) of the VSL (Figure 2a). After bonding, the VSL closed the microchannels everywhere except in the valve regions. In these locations, the through-holes in the VSL force the flow path around the obstacle in the MCL and over the valve seat, which sits 50 μm above the flexible TPU membrane (see Figure 1d). To eliminate areas of unwanted bonding, we designed this architecture, with the PMMA VSL closing the PMMA channels instead of TPU, to minimize the area where TPU bonding was necessary; we found that the bond between PMMA layers was much more consistent than the bond between PMMA and TPU since TPU could sometimes bond to the inner channel regions. Next, we used a thermal vacuum bonding technique to sandwich TPU between the valve seat side of the VSL/MCL piece and the open channel side of the PCL (Figure 2a). The resulting 125.625 cm x 92.50 cm microfluidic chip had 384 traps and 96 pneumatically controlled valves (Figure 2c,d). To complete device assembly, we attached the loading frame to the top of the MCL via pressure-sensitive adhesive (PSA) then added the outlets by gluing tubes into place.

### Valve concept and performance

We integrated microvalves to prevent drug cross-reaction between cuboids in dissimilar wells. Figure 3a shows a 3-dimensional view of the microvalve architecture, which consists of 3 main components. The primary component is the layer of TPU, a thermoplastic elastomer, whose deflection actuates the valve. The second component is a circular chamber in the control layer below the TPU that controls deflection with air pressurization. The third valve component is the valve seat. Negative pressure causes the TPU to deflect downward into the chamber, while positive pressure pushes the TPU upwards to the valve seat and blocks fluid flow. Thus, we made the shape of the valve seat half-cylindrical both to provide a path for flow when the TPU lays flat at rest, and to promote a sealing conformation with the TPU when closing the valve with positive pressure. We chose a valve seat depth of 50 μm to minimize the deflection distance required by TPU for closure, thereby reducing the required control pressure as well. Taken together, these components resulted in a normally-open, pneumatically-driven valve that could be easily fabricated across the large area of a 96-well plate using thermal vacuum bonding and operated at low pressures of ~3 psi.

**Figure 3:**
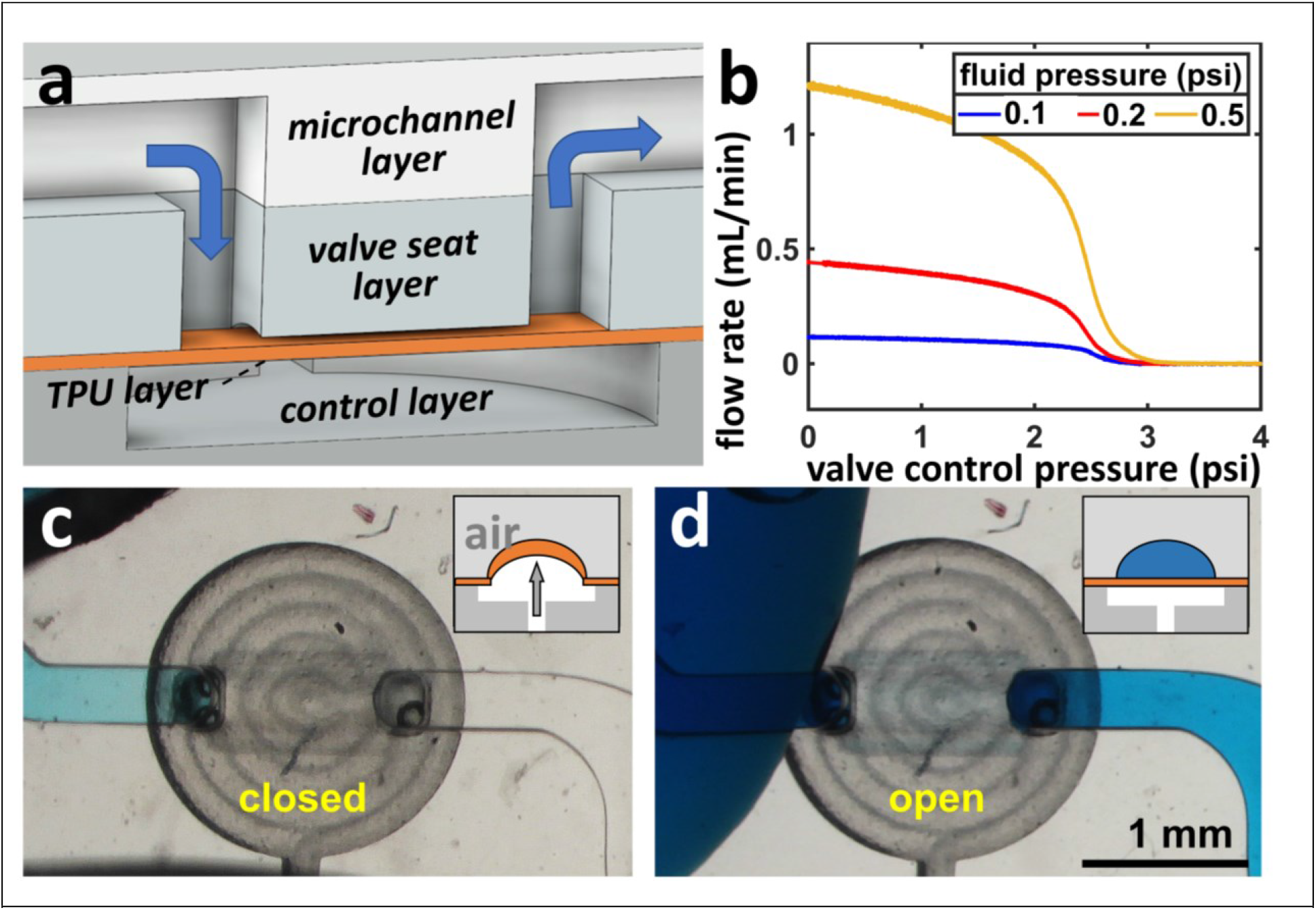
Valve design and operation. (a) 3D CAD drawing of the valve design showing the valve in the default open state. (b) Graph of flow rate versus valve control pressure (an increase closes the valve) demonstrates the effective closing of the valve at 3 different fluid pressures. (c,d) Images show how the closed valve on the device prevented flow of blue dye due to static pressure (c), and allowed flow when open (d). The insets show cross-sectional diagrams of the valve function.

Next, we evaluated valve performance. Using devices with single valves (Figure 3b), we applied constant fluid pressures (0.1, 0.2, and 0.5 psi) through the fluid channel and measured the flow rate in response to increasing valve control pressure applied through a pressure control channel. For each applied fluid pressure, +3.3 psi control pressure was sufficient to deflect TPU into the valve seat and achieve closure. In the context of our device, the valves prevented leakage at flow rates much higher than those that would be created by hydrostatic pressure. To demonstrate prevention of hydrostatic flow, we filled the fluid channels of our multi-well device with water and closed the valves with +4 psi. After we positioned wells over the traps, we filled them with blue dye to test whether the dye would leak past the valve (Figure 3c). After 40 min, we observed no leakage, and the blue dye diffused only up to the point of closure. When we opened the valves, blue dye filled the rest of the channel within 10 min (Figure 3d).

### Device operation

With the completed device, we utilized the hydrodynamic traps to array over 300 live cuboids in a 96-well organization in preparation for subsequent analysis. For this procedure, we first prepared the device by filling the channels with PBS using a handheld syringe. The application of strong positive pressure from the syringe helped to push bubbles out of traps that could otherwise prevent cuboids from entering. We connected the fluid line for one quadrant of the fluidic network to a syringe pump, and the pressure control line for the valve network to a digital pressure controller.

The cuboid capturing process is shown in Figure 4. After a rinse with PBS, we filled the central reservoir of the device with 20% PEG/PBS. We used a small PMMA frame to create a cuboid loading window within the reservoir (Figure 4b-e). As the hydrodynamic range of influence emitted by the traps was limited, the loading window circumvented this problem by congregating cuboids near the traps, increasing the likelihood that one would localize to their effective range. With negative flow generated by a syringe pump (200 mL/h total; ~1 mL/h per trap), we slid the loading window containing cuboids across the surface of the device, allowing hydrodynamic suction to pull cuboids into traps (Figure 4b,c). Once captured, cuboids remained in their trapped positions (Figure 4d,e). When valves were present, we opened the valves with −4 psi in order to minimize fluid resistance. Following this process, we repeated the process for each quadrant of the device. Separation of the trapping networks into quadrants provided a balance between simplicity of loading and the necessary flow rate. The flow rate must be sufficiently large to trap but, if made too large, it consumes a lot of PEG solution.

**Figure 4:**
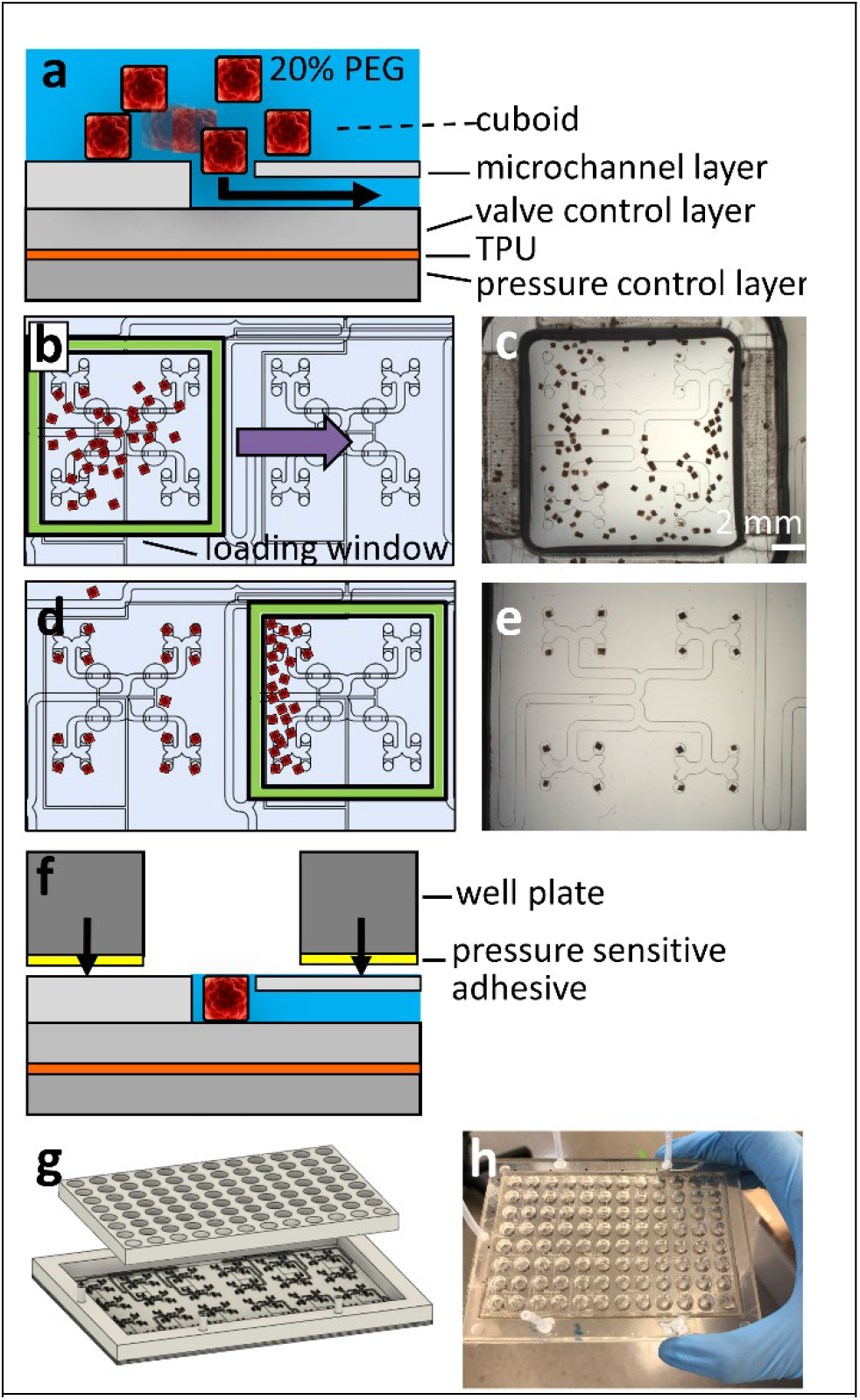
Cuboid loading. (a) Schematic shows how flow generated by suction in the outlet draws cuboids (suspended in 20% PEG) into the traps (b-e) Schematic (b,d) and images (c,e; device without valves) show how manipulation of a loading frame enables movement of the cuboids over the traps to enhance their loading. (f) Schematic of well plate layer application onto cleaned and cuboid-loaded device using pressure-sensitive adhesive. (g) 3D CAD drawing of well plate layer and device. (h) Photo of fully assembled device.

We next prepared the filled device for culture. After we dried the surface of the device via aspiration, we attached a bottomless 96-well plate to the top of the MCL with pressure-sensitive adhesive and manual pressure (Figure 4f,g). Before adding solution to the wells, we applied +4 psi of pressure to the valve control inlet to close the valves and isolate the solutions between the wells. Figure 5h shows a device after loading with live cuboids.

### Fluorescent staining of captured cuboids

To evaluate whether the valves prevented crosstalk between well conditions, we performed fluorescent staining of live cuboids in ~60 wells. We applied a repeating quad pattern of Cell Tracker Orange (CTO), Cell Tracker Green (CTG), Hoechst, or dual CTO+CTG to 4 adjacent wells in close fluidic connection with each other (Figure 6). After application of +4 psi to close the valves, we applied staining solution to each well and placed the device in a cell culture incubator for 1.5 hrs, maintaining positive pressure to the valve control inlet. Next, we imaged the cuboids after replacing the staining solutions with Live Cell Imaging Solution to stop labeling and maintain culture without the fluorescent pH indicator phenol red. Since the wells no longer required separation when filled with the same imaging solution, we disconnected the device from pressure allowing the valves to reopen. A brightfield image of the complete 96-well device with an overlay of the fluorescent cuboids (Figure 6a) indicates successfully staining of living cuboids. The same fluorescent image without brightfield highlights the differential labeling (Figure 6b). Had the valves leaked, we would have expected to see unintended staining in adjacent wells. However, cuboids in each well maintained the expected pattern of fluorescence despite sharing fluidic channels. These results indicated adequate valve closure for the duration of the assay for 75% of the wells, successfully preventing crosstalk between well conditions. These results support the potential of our 96-well-based device to perform multi-treatment assays on live tumor tissue with high throughput.

**Figure 6:**
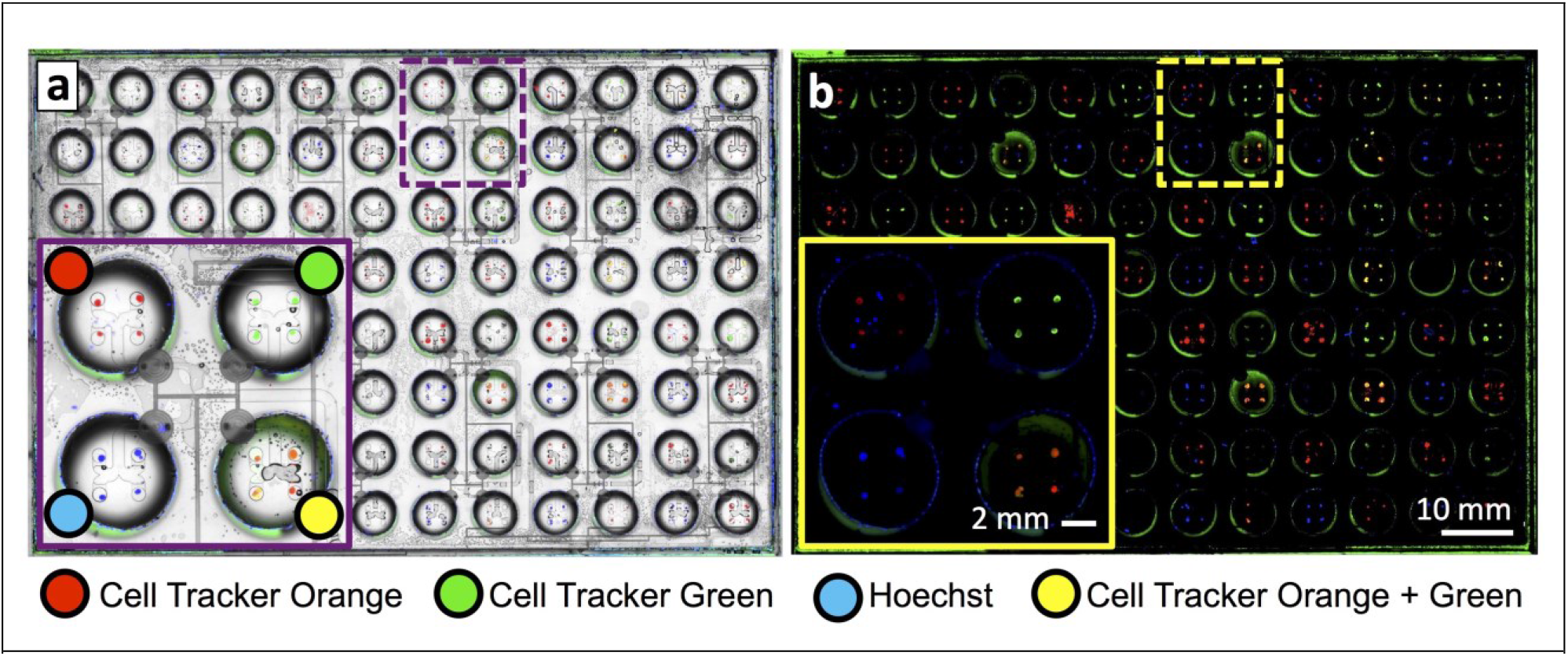
Selective fluorescent labeling of live cuboids on the microfluidic cuboid device. (a) Brightfield image overlayed with fluorescent images, and (b) fluorescent images only, for cuboids loaded into the microfluidic device, followed by labeling for one hour with different combinations of fluorescent labels whiles the valves were closed. A repeated pattern of 4 wells with different labels (as in insets) was applied across the plate, except for the lower left-hand quadrant which was not loaded with cuboids. Minimal cross-contamination was detected. Note the small blue particles in the Cell Tracker Orange well in the inset correspond to dust and not to cuboid tissue.

## CONCLUSIONS

High-throughput microfluidic approaches can empower the application of functional drug testing to microdissected tumor cuboids that preserve the tumor microenvironment. Here we presented a PMMA 96-well valved microfluidic device that simplifies the organization of hundreds of submillimeter-sized samples of tumor tissue in defined arrays for application of multiple different treatments. We trapped live cuboids from a syngeneic mouse model in a 96-well plate containing 384 traps and performed proof-of-concept dye labeling. We also showed the scalability of our manufacturing process to produce 96 completely thermoplastic microvalves that closed using low positive pressures (<4 psi), a design which may prove useful for other applications. Future, more sophisticated studies involving high numbers of cuboids could provide insights into tumor biology. Unlike our platform, present approaches for intact tissue testing have been limited thus far by low-throughput methods.

## Notes

### Competing Interest Statement

The authors have declared no competing interest.

